# Metabolomic signatures support the diagnostics of peritoneal endometriosis using generalised linear models

**DOI:** 10.64898/2026.07.05.736551

**Authors:** Alexander Cecil, Katja Vouk, Maja Novak Pušić, Andrej Vogler, René Wenzl, Cornelia Prehn, Jerzy Adamski, Tea Lanišnik Rižner

## Abstract

Endometriosis, a common inflammatory gynecological disorder affecting up to 10% of women worldwide, is characterized by the presence of endometrium-like tissue outside the uterus. Current diagnostic methods, such as ultrasound and MRI, effectively detect ovarian and deep endometriosis but fail to detect more common peritoneal type. Diagnosing peritoneal endometriosis currently necessitates invasive laparoscopy and histological confirmation. Despite numerous efforts, no new reliable biomarkers have successfully transitioned into routine clinical use.

This study aimed to investigate the use of targeted metabolomics to discover metabolite ratios capable of identifying endometriosis in plasma samples. We analyzed a discovery population of 235 patients and a validation population of 278 patients. All cases and controls in both populations were diagnosed by laparoscopy. Control subjects included individuals presenting with symptoms such as pain, dysmenorrhea, infertility, or other benign conditions, but who had no laparoscopic evidence of endometriosis. Using generalized linear models (GLMs) and machine learning, the study identified specific metabolite ratios as potential biomarkers that can distinguish different types of endometriosis and enable mass spectrometry-based diagnostics for peritoneal endometriosis. The best-validated GLM, derived from the concentration ratios of amino acids, acylcarnitines, sphingomyelins, and phosphatidylcholines, consisted of Thr/SM(OH) C22:2 + PC aa C40:5/SFA_PC + lysoPC a C16:0/SM(OH) C16:1. This model yielded an AUC of 0.82 (95% CI 0.619-0.891, with 76% sensitivity and 81% specificity) for peritoneal endometriosis. This innovative approach offers a robust diagnostic model, addressing an unmet medical need by facilitating earlier detection of peritoneal endometriosis and improving overall clinical management.

## Introduction

Endometriosis is a complex gynaecological disease characterized by endometrium-like tissue outside the uterus (ICD-10 N80 [1]). It is a benign inflammatory disease manifested by ectopic lesions on the ovaries, fallopian tubes, the tissue lining the pelvis, the bowel or the bladder [2]. Endometriosis is associated with debilitating menstrual pain, chronic pelvic pain, dysmenorrhoea, dyspareunia, dyschezia and infertility [3, 4]. The prevalence of endometriosis is slowly increasing and currently affects estimated 190 million girls and women of reproductive age or 10% of women worldwide [2, 5–7].

Despite numerous GWAS, the prevalence of endometriosis is not explained by genetics [8–10], however several risk loci were identified in multi-ancestry study [11]. The GWAS linked the pathogenesis of endometriosis to loci for infertility, heavy menstrual bleeding, cell differentiation, tissue remodelling, hormonal signal transduction pathways, and immunological processes. There are different hypotheses on the origin of endometriosis where retrograde menstruation is the prevailing but it can explain mainly peritoneal endometriosis [12, 13]. Endometriosis may further be caused by somatic mutations or epigenetic changes in endometrial stem cells [14] that cause the subsequent growth of polarised glandular structures [15]. Due to clonal selection processes within stem cell populations, metaplastic dysplasia, Muelerian rests, endometriosis exhibits different pathophysiology and manifests as ovarian, peritoneal and deep types as well as a mixed phenotype comprising combinations of those [2, 5–7].

Whilst ovarian and deep endometriosis can be detected by ultrasound or MRI scans [13], the most common peritoneal form [16] cannot be diagnosed by imaging techniques. The gold standard for the diagnosis of peritoneal endometriosis is invasive laparoscopy in conjunction with histological examination [17]. Although there is currently no cure for endometriosis, treatment is possible and should be performed as soon as possible to avoid complications such as the formation of adhesions associated with chronic pain, central sensitisation, infertility, systemic inflammation or the development of ovarian cancer [18–21]. In addition, a manifested endometriosis causes a loss of productivity and quality of life and increase sociomedical burden [22–24]. Current treatment includes hormonal therapies (progestins, GnRH agonists/antagonists and aromatase inhibitors) or surgical removal of ectopic lesions and adhesions [4, 25–28]. The decision in favour of surgery must be supported by non-invasive precision diagnostics, which are still not available. Until now, the search for individual biomarkers that enable a reliable diagnosis of endometriosis has been mostly unsuccessful. The reasons for this were incorrect study design (e.g. the use of healthy women as controls or the lack of stratification for different types of endometriosis), low specificity and sensitivity.

In the last decade, various “omics” approaches have uncovered a variety of biomarker candidates for the diagnosis of endometriosis [29–34]. Recent developments include miRNA sequencing [35–37], epigenetic analyses [38], proteomics [33], combined proteomics and metabolomics [39], nanotechnology [40] and gastrointestinal myoelectrical activity [41]. Numerous metabolomics studies have been performed to identify biomarkers for endometriosis in biological fluids such as serum, plasma, urine and peritoneal fluid [29, 39, 42]. These studies show that the pathophysiological mechanisms underlying endometriosis are associated with distinct changes in the metabolomic signatures. These changes are quantifiable and may help to overcome current diagnostic limitations. In our previous research, we performed the first targeted metabolomic analysis aimed at identifying potential biomarkers for ovarian endometriosis in both plasma and peritoneal fluid samples [43, 44]. In recent years, more studies have appeared in this area [45–49], but several either reported non-significant differences between cases and controls [45], were limited to advanced stages of endometriosis and single metabolites [46, 49] or have not validated the results [39]. A further common problem with some of these studies is that the control patients are healthy individuals. The inclusion of these individuals in the studies results of identification of biomarkers which are not endometriosis-specific but rather related to other parameters like inflammation or medication. Therefore, patients with symptoms indicative of endometriosis, including infertility and pelvic pain, but who have been found not to have endometriosis by laparoscopy should be included as controls [50]. In our recent study, we combined the protein biomarker TGFBI and metabolomics to explore logistic regression (LR) models applicable for triage testing and diagnosis of peritoneal endometriosis [51]. We identified and validated models which met the criteria for rule-in and rule-out tests and benefited from the appropriate selection of control group.

Endometriosis is a complex disease. Therefore, we hypothesized that individual metabolites would not have sufficient diagnostic value and focused instead on panels of metabolites, including metabolite ratios, which constitute metabolomic signatures. Metabolomic signatures have been shown to normalise human variability between individuals and have yielded significantly higher p and beta values in other large cohort studies of complex human phenotypes [43, 52, 53].

In this study, we investigated the potential of metabolomics to identify signatures comprising metabolite ratios characteristic of endometriosis in human plasma. The goal was to identify sets of metabolites that could be translated into a robust assay applicable for certified clinical use. Our results validated and enabled the diagnosis of endometriosis using selected metabolite ratios defined in generalized linear model (GLM) analyses. We show that GLMs can support the detection of peritoneal endometriosis, addressing an unmet medical need. The metabolites and metabolite ratios that constitute the GLMs can be reliably quantified by mass spectrometry and, after processing with machine learning techniques, provide a novel diagnostic prototype for endometriosis analysis in human blood. In this manuscript, we explain the rationale, details, advantages, and limitations of our diagnostic models.

## Methods

### Metabolomic analyses

The metabolites were quantified using flow injection and liquid chromatography-electrospray ionisation tandem mass spectrometry (FIA- and LC-ESI-MS/MS) with the AbsoluteIDQ p150 (FIA only) and p180 (FIA and LC) assays (biocrates life sciences ag, Innsbruck, Austria). The two assays allow the simultaneous quantification of 163 and 188 metabolites, respectively. Metabolites were measured from 10 µL of human plasma. The AbsoluteIDQ p180 kit covers more metabolites than the p150 kit, but both use the same technology platform and the same internal standards as calibrators for quantifying metabolites included in both assays [54, 55].

The complete test procedures and the detailed nomenclature of the metabolites have already been published [55–57]. Sample handling was performed using a Hamilton Microlab STAR robot (Hamilton Bonaduz AG, Bonaduz, Switzerland) and an Ultravap nitrogen vaporiser (Porvair Sciences, Leatherhead, UK), in addition to standard laboratory equipment. In brief, 10 µL of plasma was added to the cavities of the 96-well filter plate of the p150 or p180 assay and dried in a nitrogen stream for 30 minutes. Amino acids and biogenic amines were derivatised with an excess of 5% phenylisothiocyanate in ethanol/water/pyridine (1/1/1 v/v/v) for 20 minutes and then dried. Samples were extracted for 30 minutes at room temperature with 300 µL methanol containing 5 mM ammonium acetate and diluted differently for the FIA and LC runs. The LC run was performed with an Agilent XDB-C18 column (3 × 100 mm, 3.5 µm). Mass spectrometric analyses were performed using an API4000 triple quadrupole system (SCIEX Deutschland GmbH, Darmstadt, Germany) equipped with a 1260 series HPLC (Agilent Technologies Deutschland GmbH, Böblingen, Germany) and an HTC-xc PAL autosampler (CTC Analytics, Zwingen, Switzerland), controlled by Analyst 1.6.2 software. For the LC-MS/MS part, compounds were identified and quantified based on scheduled multiple reaction monitoring measurements (sMRM); for the FIA-MS/MS part, on the basis of MRM. Data analyses for the quantification of metabolite concentrations and quality assessment were performed using MultiQuant 3.0.1 software (SCIEX) and the MetIDQ software package, which is an integral part of the AbsoluteIDQ kit. Metabolite concentrations were calculated using internal standards and expressed in µmol/L (µM).

Measurements were performed at the metabolomics platform of the Genome Analysis Centre at Helmholtz Zentrum München. The discovery and validation populations were measured in separate batches using AbsoluteIDQ p180 and p150 kits, respectively. In addition to the study samples, three quality controls provided by the manufacturer, three zero samples (solvents only), and a further five aliquots of pooled human reference plasma were analysed on each kit plate. The results of the reference plasma aliquots were used to calculate possible batch effects and to normalise the data. After quality control (QC), metabolites were retained if the average coefficient of variation (CV) in the reference or QC samples was <25 and if more than 50% of the measured concentrations were above the limit of detection (LOD), defined as three times the median of the zero samples. If the analytes in a given sample were below the LOD, “NA - not available” was used in the dataset.

Metabolite abbreviations follow the nomenclature as described in [58]. Specifically, acylcarnitines (e.g. C6:1) are denoted as x:y, where x and y represent the total number of carbons and double bonds in all chains, respectively. Amino acids are abbreviated with three letters (e.g. Gln). Phosphatidylcholines are indicated as PC nn x:y, where nn is either “aa” for diacyl or “ae” for acyl-alkyl. Lysophosphatidylcholines are denoted as lysoPC a x:y, where “a” stands for acyl. Sphingolipids are indicated as SM x:y. “na” indicates not annotated in the HMDB (Human Metabolome Data Bank) [59]. The chemical names of the metabolites are given without a resolution of the isobars. The names of the individual metabolites are defined as state of the art in the research field [43, 44, 60]. Further metabolite characteristics such as trivial names, CAS numbers and HMDB accession numbers can be found in the supplementary Table S1.

The primary metabolomic dataset from the two study populations was recently analyzed using the LR approach and combined with protein data (concentrations of TGFBI) [51].

### Analysis populations

Human plasma samples were collected in accordance with standard operating procedures (SOPs) [61] at the Departments of Obstetrics and Gynaecology, University Medical Centre Ljubljana, Slovenia, and the Medical University of Vienna, Austria, from March 2009 to February 2016 for the validation population, and at the University Medical Centre Ljubljana, Slovenia, from March 2016 to January 2019 for the discovery population. A detailed demographic description of the cohorts is provided in supplementary Table S2. The study was conducted in accordance with the Declaration of Helsinki and was approved by both national medical ethics committees (Slovenian approval no. 110/05/14, and no. 49/03/16; Austrian approval no. 545/2010). All participants provided written informed consent before participating in the study.

Additional information about the patients’ lifestyle, gynaecological and clinical status was collected. The patients completed a comprehensive questionnaire developed by our research group [43] that assessed their medical history, stress levels, medication use, diet, lifestyle habits, type of pain (dyspareunia or chronic pain), and dysmenorrhoea, using a validated visual analogue scale [62]. Other clinical information and additional pathologies were documented by the surgeon performing the procedure using a separate questionnaire. This questionnaire included details on the length and regularity of the menstrual cycle, use of oral contraceptives and/or hormone therapy in the past and in the three months prior to surgery, medication taken one week before surgery, type and reason for surgery, type of endometriosis, revised American Society for Reproductive Medicine (rASRM) stage, colour of lesion, histological confirmation of endometriosis, menstrual phase determined, and additional pathological observations. Disease status (case or control) was confirmed by laparoscopic surgery. Control subjects were defined as those with phenotypes such as pain, dysmenorrhoea, infertility, or other benign conditions, but who were not diagnosed with endometriosis by laparoscopy. Exclusion criteria included pregnancy, age under 18 or over 50 years, confirmed menopausal status, gynaecological malignancies, cancelled surgery, previous hysterectomy, missing clinical data, and other comorbidities. All patients were of European origin. Samples were stored at −80 °C until analysis. Measurements were performed in May 2020 and June 2016 for the discovery and validation populations, respectively.

The analysis populations were classified by health status and endometriosis type, with subsequent adjustment for body mass index (BMI) and age. Adult BMI categories were defined according to standard NIH/WHO cut-offs: underweight (<18.5), normal weight (18.5–24.9), overweight (25.0–29.9), and obesity (≥30.0) [63, 64]. Patients with a BMI ≥ 30 (9 cases, 5.9% (9/151) of all cases; 11 controls, 13.09% (11/84) of all controls), corresponding to 8.5% (20/235) of the entire discovery population, were not excluded from the study. This approach allowed for the assessment of natural distribution and BMI as a potential confounder.

Figure 1 presents an overview of the sample distribution after applying the exclusion criteria to the control and case groups across different populations, along with a schematic representation of the overall study design. The cases are separated into peritoneal, ovarian, and deep endometriosis (Figure 1). Cases with multiple mixed types of endometriosis are included in the ‘mixed’ category. Specifically, the “ovarian mixed” group comprises patients with both ovarian and peritoneal types, the “peritoneal mixed” group consists of patients with both peritoneal and deep types, and the “deep mixed” group includes patients with deep, ovarian, and peritoneal types of endometriosis.

**Figure 1.**
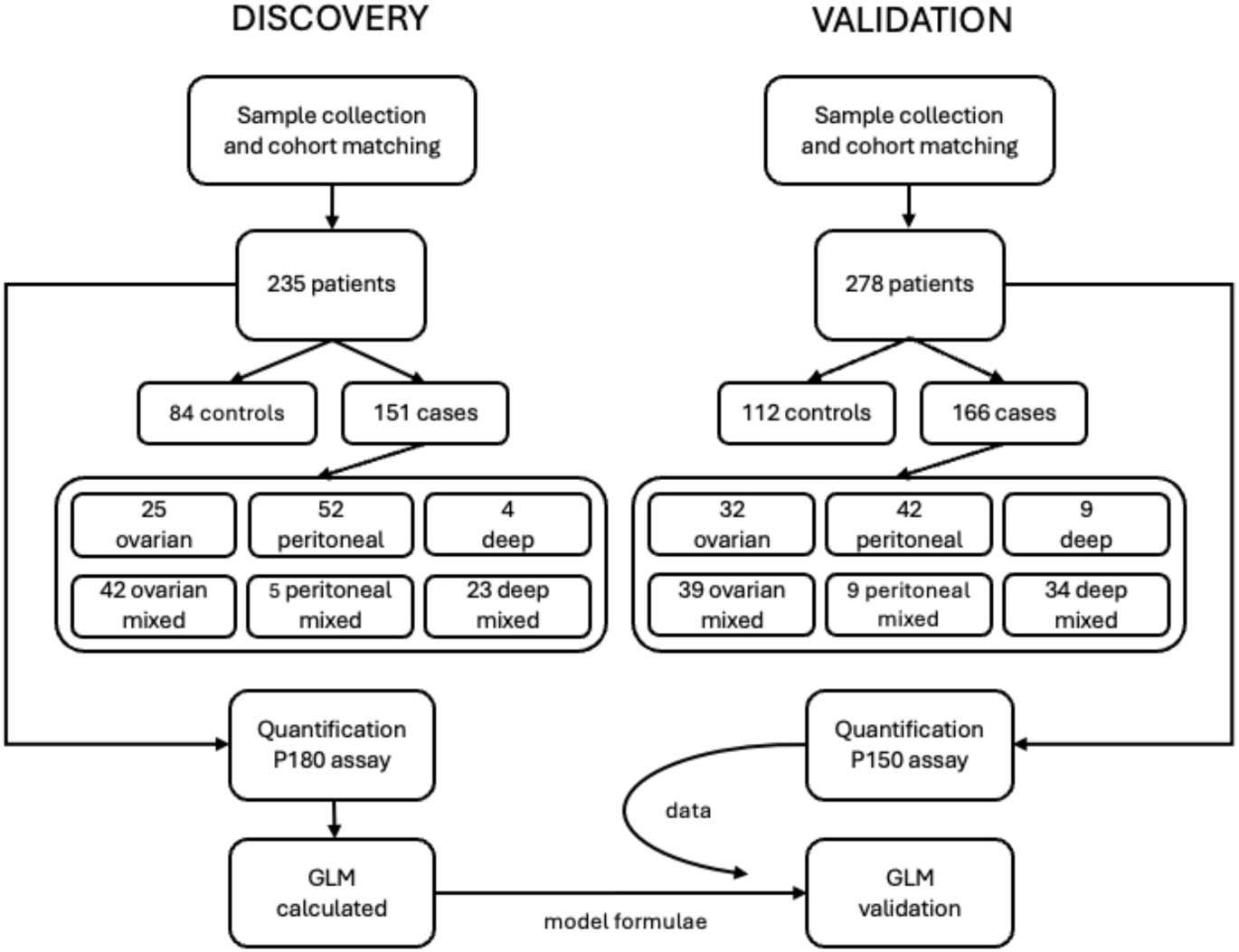
Overall flow and distribution of phenotypes in the discovery and validation cohorts. The number of individual patients is indicated for each phenotype.

### Statistics

A power calculation [65] was performed based on the assumption of a target specificity of at least 85% and a minimum sensitivity of 90%, resulting in a requirement of at least 138 cases.

Only metabolites present in both the AbsoluteIDQ p180 and p150 datasets were processed further. The measurement results from both the discovery and validation populations were processed using the same statistics method.

Statistical analyses were performed with R (version 4.0.5) [66]. R scripts were used to detect missing data and for general quality control of metabolites and samples. Detection of NA values was performed, and the metabolites OH-Pro, carnosine, DOPA, dopamine, nitro-Tyr, serotonin, and tryptophan were excluded as they had more than 40% NA across all samples and were therefore considered too unreliable.

Imputation was performed for metabolites with less than 40% below LOD/LLOQ i.e. for NA. As the minimum detectable value is recorded for each metabolite, this value was then divided by the square root of 2 [67]. A random permutation of ±25% was applied around this recalculated imputation value to mitigate biassed statistical results and to avoid the presence of identical values after imputation.

Further CV tests were also performed and C16:1-OH, C3-OH, histamine, phenyl-ethylamine, symmetric dimethylarginine, serotonin, spermine, spermidine, PC aa C24:0, PC aa C30:2, PC ae C30:1, lysoPC a C26:0, lysoPC a C26:1, lysoPC a C28:0 and lysoPC a C28:1 were removed due to a coefficient of variation (CV) greater than 25%.

The following workflow was used to analyse the data. The dataset was divided into five separate analysis treads: 1) endometriosis stages within the same endometriosis type were not distinguished to obtain a very general model; 2) all endometriosis types and controls; 3) the deep type was filtered out due to the small number of samples; 4) ovarian and control samples; 5) ovarian, the mixed ovarian-peritoneal type, and control samples; 6) peritoneal and control samples; 7) peritoneal, the mixed ovarian-peritoneal type, and control samples. For all these datasets, all possible metabolite ratios were independently calculated and appended to the datasets.

Due to the inherent left shift of metabolomics data, which in most cases prohibits normal data distribution even after log transformation and/or Pareto scaling, Shapiro-Wilk testing was performed to elucidate data distribution. It indicated non-normal data distribution in at least 30% of cases. This threshold was deemed too high to assume normal data distribution and proceed with parametric statistical testing [68–70]. Therefore, the Wilcoxon rank-sum test was chosen to evaluate statistical significance. To that end, the ‘wilcox.test’ function [71, 72] in R (version 4.0.2 (2020-06-22) [66] was applied to the data. However, this classical statistical approach proved insufficiently stringent for the samples included. Therefore, the final selection of metabolites from non-log-transformed datasets was aided by machine learning, specifically randomForest /RF) (number of trees = 2000) [73]. The input data for randomForest was based on the statistically significant results of all metabolites and all possible metabolite ratios. This pre-selection of potential biomarker targets was always performed with a p-value of 0.05, without correction for multiple testing.

The machine learning algorithm chosen for this task was the ‘randomForest’ package from R (version 4.7-1.1)[73]. The number of trees was set to 2000, and all calculations were performed on 10-fold cross-validated data. For each cross-validation step, the samples from the discovery datasets were randomly divided into two-thirds training data and one-third test data, with only the training data used to calculate the random forest models. For each cross-validation step, only the results of these random forest calculations that ranked in the top 10% according to the “MeanDecreaseGini” performance criterion were selected. Of all possible candidates from the 10-fold cross-validation steps, only the overall best 10% were considered for further modelling.

These further prediction models were created using generalised linear models (‘GLM’, R package ‘glm’ version 4.2.1 [74–77]. For these models, following the PLS-R predictions, we decided to use 3 predictors to create all possible models. The calculations of confidence intervals (CI) for each AUC were performed using the bootstrap method (2000 bootstraps)) [78].

## Results

### The metabolomic signatures of the controls and cases are similar but not identical

In the discovery population, we first analysed the distribution of metabolite signatures using partial least squares-discriminant analysis (PLS-DA) to assess differences between case and control groups. The groups are not identical, but there is substantial overlap, as shown in Figure 2.

**Figure 2:**
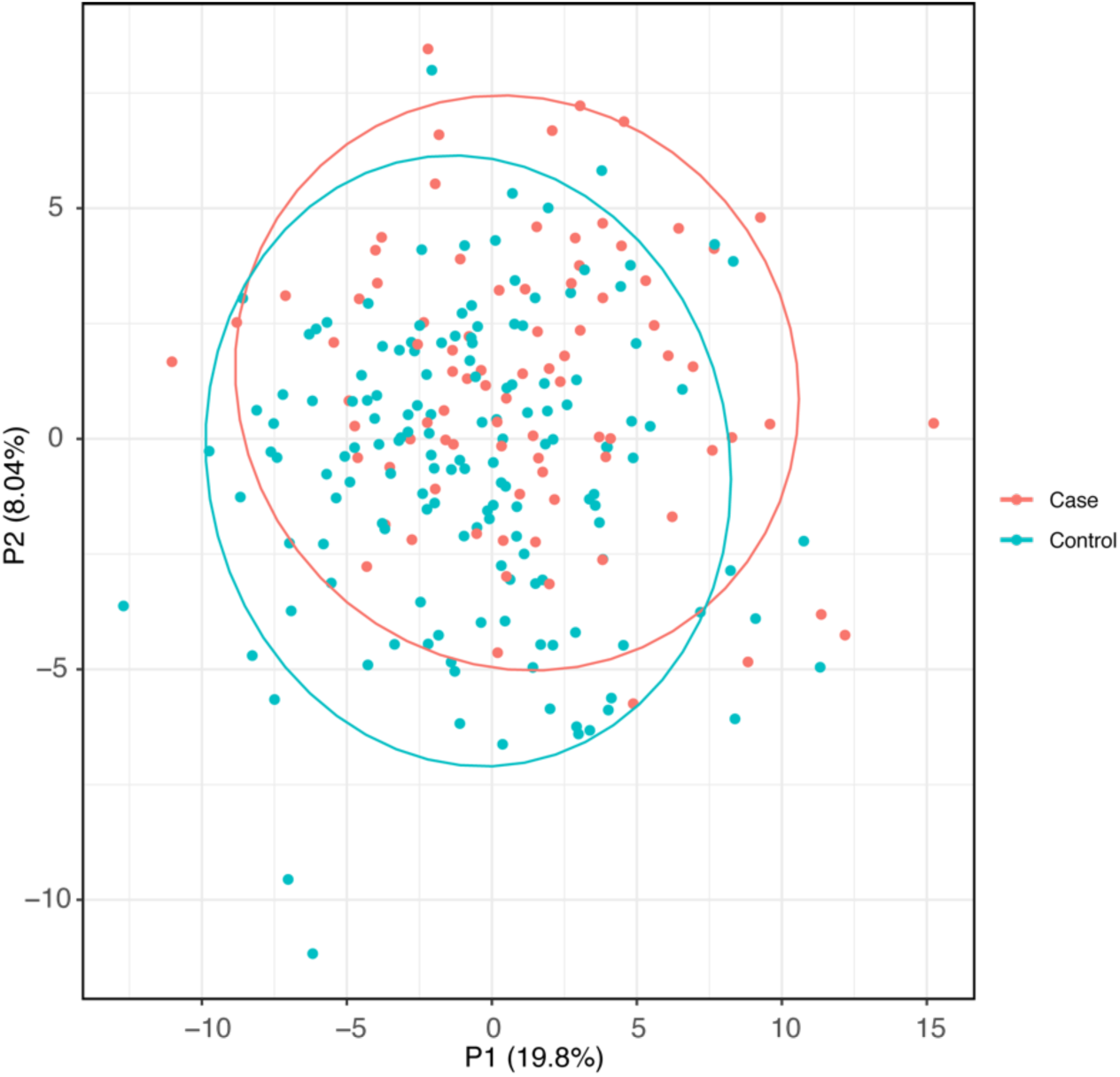
Analysis of differences in metabolite signatures between endometriosis patients and controls using PLS-DA does not support differentiation between cases and controls. In this figure, cases include all types of endometriosis. The dataset comprised controls (n = 151) and cases (n = 84).

The data used in the Figure 2 were non-log transformed metabolite concentrations. This PLS-DA does not include the metabolites which were found to be above the CV% threshold of 25% or were excluded due to being above the NA threshold of 40%. As is evident from the PLS-DA statistics, a separation by group (i.e. control vs case) is not possible if based on absolute concentrations of metabolites. The statistic parameters of PLS-DA in Figure 2 are as follows: R*_γ_*^2^ = 0.326; R^2^_X_= 0.47; Q_x_^2^ = −0.422; RMSEE = 0.4; P_R_^2^ = 0.288; P_Q_^2^ = 0.3165 which illustrates a lack of distinction of case and controls as defined in [79–82]. Specifically, the parameters are defined as follows: R*_γ_*^2^ describes explained variation and should be above 0.75 and never negative; R^2^_X_ represents the variance explained predictor variables, and should be between 0.4-0.9, Q_x_^2^ describes predicted variation and should be above 0.4, never 1.0 and never negative; RMSEE is the root mean square error of estimations and describes the accuracy of the model. It should be below 0.25; P_R_^2^ is the R^2^ parameter after permutation testing (2000 times) of sample grouping which has to be below 0.05 to produce a valid PLS-DA; P_Q_^2^ depicts the Q^2^ parameter after permutation testing (2000 times) of sample grouping and has to be below 0.05 to produce a valid PLS-DA [79–82].

### Common confounders do not affect metabolomic signatures of controls and cases

Using the discovery dataset, we analysed the influence of the menstrual cycle on plasma metabolomic signatures in both controls and cases (Figure 3). The results of the PLS-DA analysis of the explanation of menstrual phase by metabolites did not show any significant dependencies, and the specific phenotypes were not separated as groups. The dendrogram calculated using the Ward algorithm [83] with a Euclidean distance measure also did not reveal any clusters (not shown).

**Figure 3.**
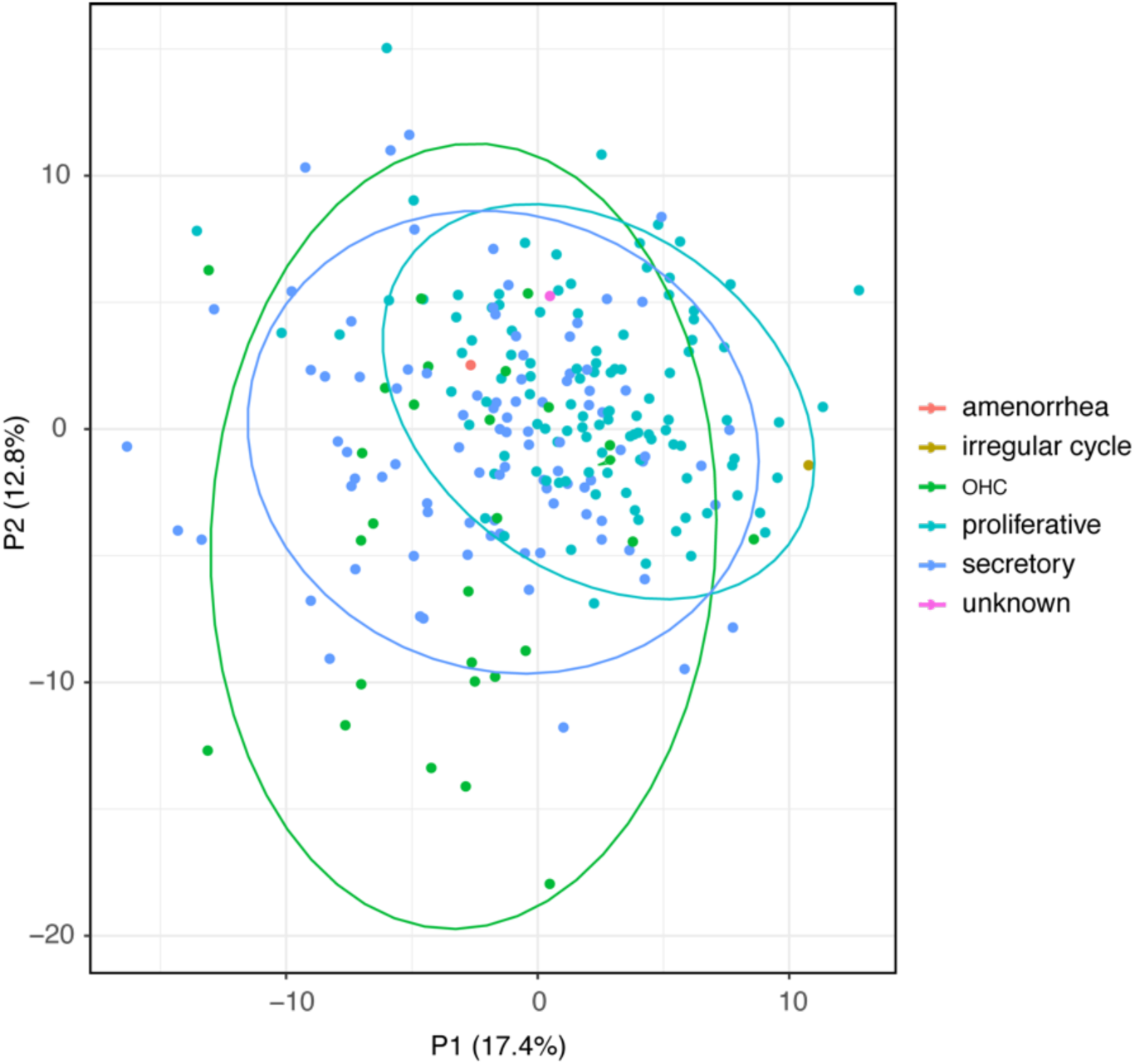
The menstrual cycle does not support stratification of patients according to the metabolomic signatures. The different phases of the menstrual cycle are indicated as follows: blue for the secretory phase (n = 94), turquoise for the proliferative phase (n = 110), green for the oral hormonal contraception (OHC) group (n = 20), beige for irregular cycle (n = 1), orange for amenorrhoea (n = 1), and pink for unknown (n = 1). The parameters for the PLS-DA are as follows: R*_γ_*^2^ = 0.185; R^2^_X_= 0.488; Q_x_^2^ = −0.0277; RMSEE= 0.266; P_R_^2^ = 5e^−4^; P_Q_^2^ = 5e^−4^.

Other confounding factors tested, such as age, BMI (even after inclusion of patients with BMI over 30), contraception, smoking habits, medication and alcohol consumption, had no statistically significant influence on the distribution of metabolomics signatures (Figure 4) and did not provide a clear separation of phenotype. In other words, the confounders do not differentiate patients into clearly separated groups.

**Figure 4.**
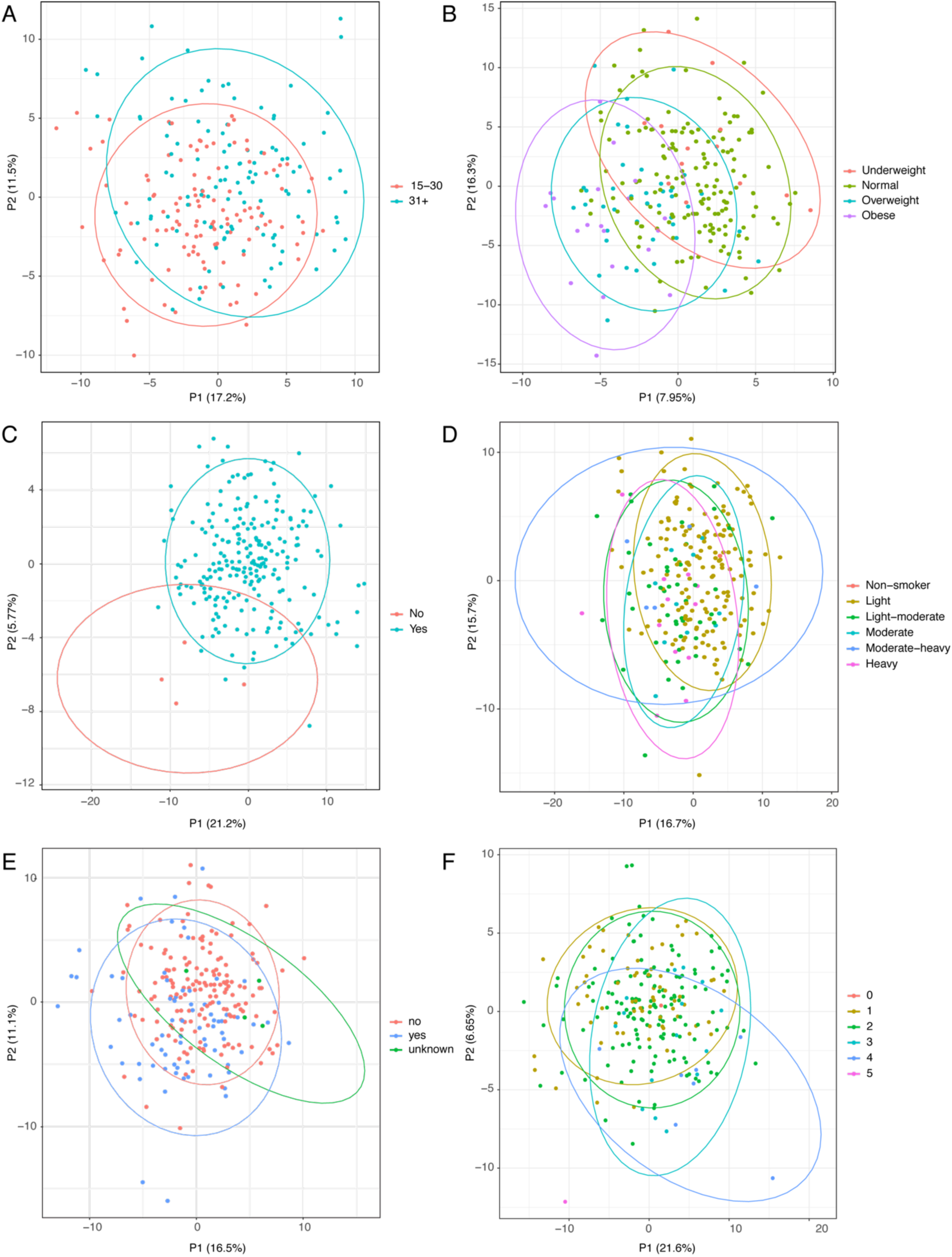
The confounding factors age, BMI, contraception, smoking, medication, and alcohol consumption do not contribute significantly to explaining the distribution of metabolomic signatures by PLS-DA. A) Age with the criteria “15-30 years old” (red, n = 120) and “31+” (green, n = 115); B) BMI labelled with the criteria “underweight” (blue, n = 13), “normal” (orange, n = 161), “overweight” (green, n = 41), and “obese” (beige, n = 20); C) Contraception “No” (turquoise, n=141), “Yes” (orange, n=9); D) Smoking with the criteria “non-smoker” (orange, n = 1), “light” (beige, n = 160), light to moderate (green, n = 47), moderate (turquoise, n = 10), moderate-heavy (blue, n = 5) and heavy (pink, n = 12), E) Medication with data for none (red, “no” n = 159), present (blue, “yes” n = 72) and unknown (green, n= 4), and F) Alcohol consumption in categories unknown (“0” orange, n = 1), never (“1” brown n =68), occasionally (“2” green, n = 143), once a week (“3” turquoise, n=15), two to 3 times a week (“4” blue, n= 7) and more than 3 times a week (“5” red, n = 1). The PLS-DA parameters for the specific confounders are: A) Age, R*_γ_*^2^ = 0.234; R^2^_X_= 0.469; Q_x_^2^ = − 0.176; RMSEE= 0.363; P_R_^2^ = 0.0665; P_Q_^2^ = 0.0765; B) BMI, R*_γ_*^2^ = 0.213; R^2^_X_= 0.478; Q_x_^2^ = − 0.00307; RMSEE= 0.28; P_R_^2^ = 5e− 4; P_Q_^2^ = 5e− 4; C) Contraception, R*_γ_*^2^ = 0.355; R^2^_X_= 0.46; Q_x_^2^ =− 0.0999; RMSEE= 0.189; P_R_^2^ =0.0475; P_Q_^2^ = 0.0045; D) Smoking, R*_γ_*^2^ =0.146; R^2^_X_= 0.474; Q_x_^2^ =− 0.0708; RMSEE= 0.262; P_R_^2^ =0.001; P_Q_^2^ = 0.0145; E) Medication, R*_γ_*^2^ =0.204; R^2^_X_= 0.471; Q_x_^2^ =− 0.0235; RMSEE= 0.331; P_R_^2^ =0.6195; P_Q_^2^ = 0.2155 and F) Alcohol, R*_γ_*^2^ =0.116; R^2^_X_= 0.474; Q_x_^2^ =− 0.159; RMSEE= 0.277; P_R_^2^ =0.314; P_Q_^2^ = 0.5485.

### Generalised Linear Models differentiate controls and cases

As the classical statistical approach proved insufficient for detecting differences in the dataset, metabolite selection was performed using machine learning with RF on all metabolites and all possible metabolite ratios. Individual metabolite ratios have been reported to contribute to the normalisation of individual variability in human parameters in various omics studies [43, 52, 53]. However, in this study we wanted to investigate models that include composite signatures containing single metabolite and/or multiple metabolite ratios.

All calculations were performed using 10-fold cross-validated data. For each cross-validation step, the data were randomly divided into 66% training data and 34% test data. To narrow down the possible candidates for further modelling with generalised linear models (GLMs) and to obtain receiver operator characteristic (ROC) curves with area under the curve (AUC) calculations, the following criteria for random forest (RF) were used: number of computed trees: 2000, MeanDecreaseGini: > 0. Of the remaining candidates, only those in the top 10% of performance were selected. We found that models containing up to three metabolite ratios and/or a single metabolite showed the greatest improvement in AUCs with moderate computational time requirements (e.g. two weeks). Models with four or more metabolite ratios increased AUCs only minimally but required much longer calculation times (e.g. months). For the remaining candidates, all possible combinations were calculated for three-ratio predictor models for the GLM. Multiple GLMs with different AUCs were calculated for each type of endometriosis. The following models were calculated: 1) all endometriosis types: 43,744 models; 2) ovarian: 23,478; 3) ovarian mixed: 43,744 (same number as for all endometriosis types, but different models); 4) peritoneal: 24,857; 5) peritoneal mixed: 45,825.

### Analysis of discovery population supports the development of models that differentiate peritoneal endometriosis from other types of disease

GLMs were calculated modeling the case-control outcome of samples in the discovery population, Below, we describe the best models for peritoneal endometriosis. Only the five best models are listed in Table 1, ranked by AUC. The identities of the metabolites in the GLMs are provided in Supplementary Table S1, and SFA_PC refers to the composite lipid class of saturated glycerophosphocholines.

**Table 1.** Different GLMs for peritoneal endometriosis in the discovery population.

| # | GLM | AUC<br>average | 95%CI | RMSE | SP<br>% | SE<br>% |
| --- | --- | --- | --- | --- | --- | --- |
| 1 | Thr/SM(OH) C22:2 + PC aa C40:5/SFA_PC + lysoPC a C16:0/SM(OH) C16:1 | 0.83 | 0.61-0.89 | 0.09681 | 76 | 81 |
| 2 | PC aa C40:5/SFA_PC + lysoPC a C16:0/SM(OH) C16:1 + PC aa C32:0/SM C18:0 | 0.81 | 0.60-0.88 | 0.08131 | 75 | 75 |
| 3 | lysoPC a C16:0/SM(OH) C16:1 + PC aa C32:0/SM C18:0 + PC aa C32:0/PC aa C38:3 | 0.80 | 0.59-0.87 | 0.08836 | 75 | 75 |
| 4 | C6:1_div_by_Pro/PC aa C40:5/SFA_PC + lysoPC a C16:0/SM(OH) C16:1 | 0.80 | 0.63-0.89 | 0.09442 | 76 | 82 |
| 5 | PC aa C40:5/SFA_PC + lysoPC a C16:0_div_by_SM(OH) C16:1 + Arg/PC ae C34:0 | 0.80 | 0.61-0.88 | 0.09739 | 75 | 72 |

Abbreviations: metabolite abbreviations are explained in supplementary Table S1, # - number of the model, AUC – area under the curve, 95%CI – range of confidence interval at 95%, RMSE - root mean square error, SP – specificity, SE – sensitivity.

The models are listed according to the average AUC and the RMSE (Root Mean Squared Error) shows less than 0.15 in all cases, which indicates the statistical validity of the models, as explained in [84, 85]. The cross-validation calculations for AUC for model #1 in Table 1 are shown in Figure 5. For the sake of clarity, only one model is shown. Other models from Table 1 are shown in Supplementary Figure S1.

**Figure 5.**
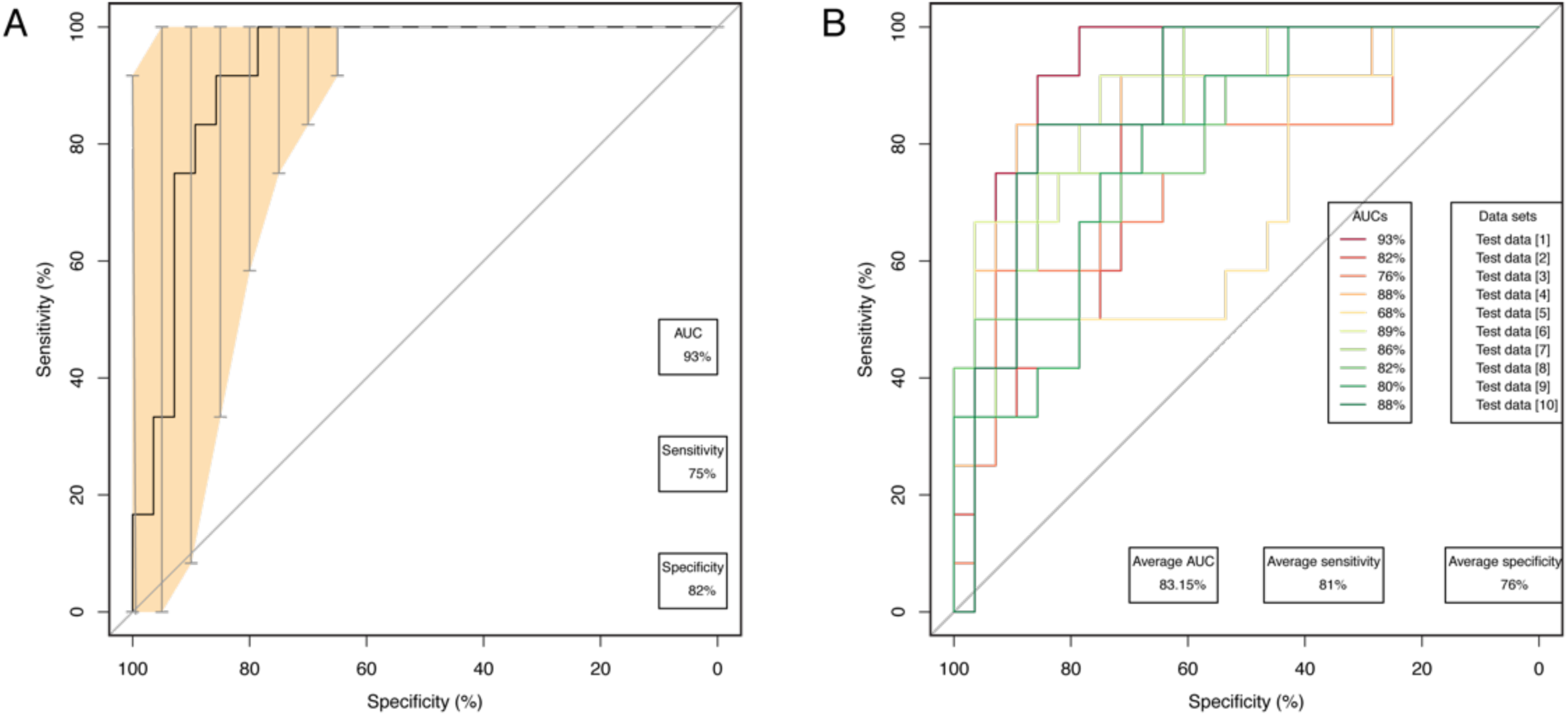
AUC parameters for the best model to discriminate peritoneal endometriosis from control group in the discovery population. Data are shown for a model containing the following metabolites: Thr/SM(OH) C22:2 + PC aa C40:5/SFA_PC + lysoPC a C16:0/SM(OH) C16:1. A) best cross-validated model, the light orange area and bars represent 95%CI, B) the model was cross-validated 10 times and the mean AUC, specificity and sensitivity, and further individual AUCs, are given.

The GLMs yielded a mean AUC of 0.83, with the RMSE below the threshold of 0.15, again indicating the statistical validity of the models.

The other types of endometriosis were also analysed, yielding the AUCs shown in Table 2. The AUCs of the GLMs for peritoneal endometriosis performed best among the types examined. No statistically valid models were calculated for cases with deep endometriosis and mixed cases of deep endometriosis, including ovarian/deep endometriosis, peritoneal/deep endometriosis, and ovarian/peritoneal/deep endometriosis, due to high variability of metabolomic signatures in comparison to the number of cases.

**Table 2.** Examples of GLMs for best diagnostic performance in different types of endometriosis in the discovery population.

| Endometriosis type | GLM formula | AUC | 95%CI | SP % | SE % |
| --- | --- | --- | --- | --- | --- |
| All types | LysoPC a C17:0/SM(OH) C16:1 + Arg/PC ae C36:0 + PC ae C38:0/PC ae C40:0 | 0.72 | 0.58-0.80 | 85 | 72 |
| Peritoneal | Thr/SM(OH) C22:2 + PC aa C40:5/SFA_PC + lysoPC a C16:0/SM(OH) C16:1 | 0.83 | 0.61-0.89 | 76 | 81 |
| Peritoneal mixed | Orn/PC ae C38:0 + C4/PC aa C38:4 + Tyr/PC aa C42:2 | 0.68 | 0.47-0.75 | 69 | 64 |
| Ovarian | PC aa C36:3/PC ae C40:5 + lysoPC a C14:0/PC aa C28:1 + Met/PC aa C36:3 | 0.70 | 0.61-0.89 | 92 | 77 |
| Ovarian mixed | C10/PC aa C36:6 + Pro/PC ae C34:0 + PC ae C42:3/SM(OH) C16:1 | 0.66 | 0.49-0.75 | 63 | 60 |

Abbreviations are explained in Table 1.

The values for AUC, specificity, and sensitivity represent the averages of the results after 10-fold cross-validation. The corresponding figures for cross-validation are shown in Supplementary Figure S2. Supplementary Table S3 also lists other GLMs for each endometriosis type that have lower AUC values.

### Validation of GLMs demonstrates the validity of observations that are relevant for diagnostics

We aimed to validate the results for peritoneal endometriosis in an independent cohort analysed using the absolute concentration quantification method for the same metabolites. Therefore, the models presented in Table 1, obtained in the discovery population, were applied to the validation population, which comprises samples from different patients than those in the discovery phase and includes an additional study centre. The validated results for the different GLMs for peritoneal endometriosis are shown in Table 3.

**Table 3.** Validated parameters of GLMs for peritoneal endometriosis.

| # | GLM formula | AUC | CI 95% | SP % | SE % |
| --- | --- | --- | --- | --- | --- |
| 1 | Thr/SM(OH) C22:2 + PC aa C40:5/SFA_PC + lysoPC a C16:0/SM(OH) C16:1 | 0.82 | 0.69-0.84 | 71 | 41 |
| 2 | PC aa C40:5/SFA_PC + lysoPC a C16:0/SM(OH) C16:1 + PC aa C32:0/SM C18:0 | 0.79 | 0.67-0.83 | 78 | 91 |
| 3 | lysoPC a C16:0/SM(OH) C16:1 + PC aa C32:0/SM C18:0 + PC aa C32:0/PC aa C38:3 | 0.80 | 0.64-0.81- | 60 | 33 |
| 4 | C6:1/Pro/PC aa C40:5/SFA_PC + lysoPC a C16:0/SM(OH) C16:1 | 0.79 | 0.71-0.86 | 80 | 94 |
| 5 | PC aa C40:5/SFA_PC + lysoPC a C16:0/SM(OH) C16:1 + Arg/PC ae C34:0 | 0.80 | 0.67-0.83- | 78 | 91 |

Abbreviations are explained in Table 1.

As indicated by the AUC values, the models were validated, albeit with varying sensitivity and specificity.

We further analysed the performance of the GLMs in other types of endometriosis. The overview of the validation results for the different types is shown in Table 4.

**Table 4.**
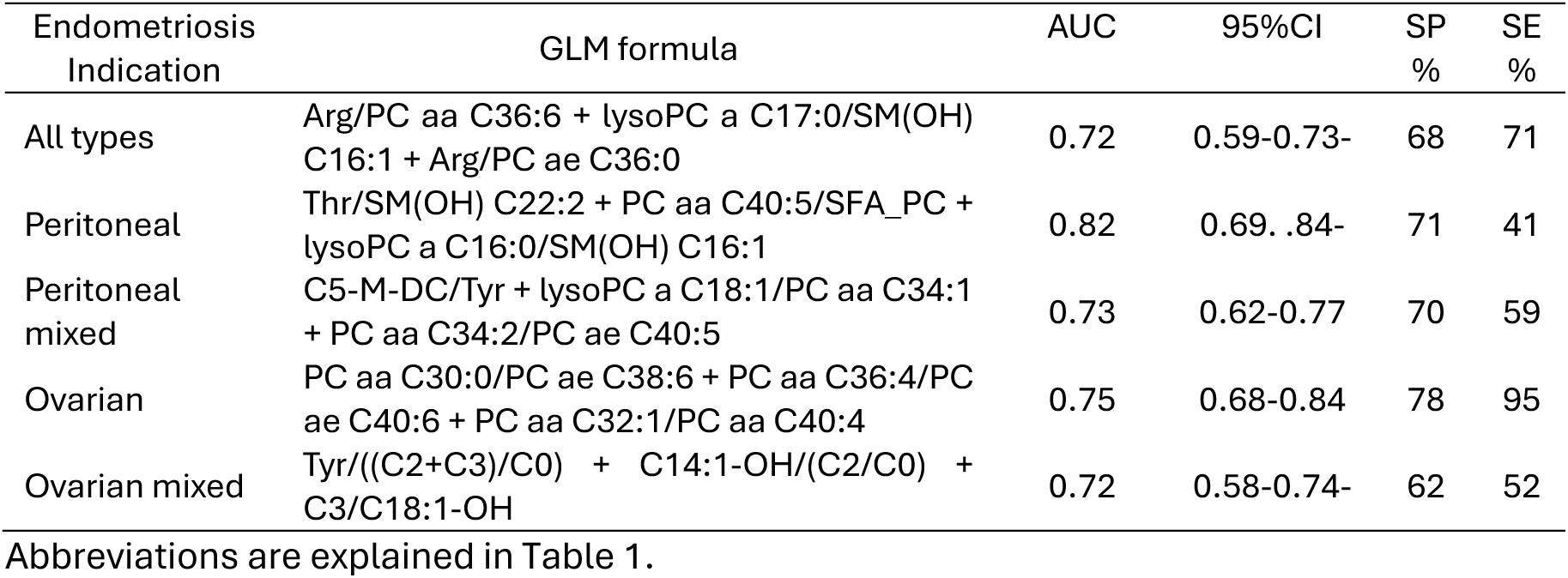
Performance of GLMs in the validation population for distinct types of endometriosis.

| Endometriosis Indication | GLM formula | AUC | 95%CI | SP % | SE % |
| --- | --- | --- | --- | --- | --- |
| All types | Arg/PC aa C36:6 + lysoPC a C17:0/SM(OH) C16:1 + Arg/PC ae C36:0 | 0.72 | 0.59-0.73- | 68 | 71 |
| Peritoneal | Thr/SM(OH) C22:2 + PC aa C40:5/SFA_PC + lysoPC a C16:0/SM(OH) C16:1 | 0.82 | 0.69-.84- | 71 | 41 |
| Peritoneal mixed | C5-M-DC/Tyr + lysoPC a C18:1/PC aa C34:1 + PC aa C34:2/PC ae C40:5 | 0.73 | 0.62-0.77 | 70 | 59 |
| Ovarian | PC aa C30:0/PC ae C38:6 + PC aa C36:4/PC ae C40:6 + PC aa C32:1/PC aa C40:4 | 0.75 | 0.68-0.84 | 78 | 95 |
| Ovarian mixed | Tyr/((C2+C3)/C0) + C14:1-OH/(C2/C0) + C3/C18:1-OH | 0.72 | 0.58-0.74- | 62 | 52 |

Abbreviations are explained in Table 1.

The discovery population models were validated in an independent population, achieving AUC values between 0.72 and 0.82. Although the exact discovery model produced a slightly lower AUC during validation, the best-performing models from the discovery cohort were also among the best-performing models in the validation cohort.

## Discussion

### Biomarkers are needed for the early diagnosis and therapy monitoring of endometriosis

There is no definitive cure for endometriosis, but early intervention with hormonal or surgical treatments is essential for efficient management of disease [18–21]. Recent research has shifted to metabolomics or combination of proteins and metabolites as biomarkers [51], focusing on metabolite ratios rather than individual metabolites to normalise human variability and improve diagnostic accuracy [42]. These metabolites and ratios, which are measurable by mass spectrometry and, as shown in this study, can be refined using machine learning, offer promising advances in non-invasive diagnostic methods.

### Other approaches in search for the biomarkers of endometriosis

Serum analyses of miR-17, IL-4, and IL-6 levels initially showed promising diagnostic potential for early endometriosis, however, their lack of specificity limits clinical applicability, as recently demonstrated [34, 86]. Proteomic profiling of 805 patients produced a 10-protein panel with an impressive AUC of 0.99; however, common confounding variables such as age and BMI were not considered, patients were compared to healthy women, and the models were not validated [33]. While the miRNA panels from plasma showed modest accuracy (AUC 0.60) [87], the miRNA signatures from saliva yielded better results (AUC 0.96) [88, 89]. This AUC might be inflated by covariates (i.e. over 100 miRNA species), which were not investigated. The individual miRNAs in this study were also altered in several other common diseases such as NAFLD, neuronal dysfunction, rheumatoid arthritis, and type 1 and type 2 diabetes [90]. It is also known that miRNA profiling is susceptible to potential contamination [91], which does not affect proteomic and metabolomic approaches. A recent combined metabolomics and proteomics study claimed to have discovered 20 metabolites relevant for peritoneal fluid diagnostics and 26 metabolites relevant for plasma-based diagnostics of endometriosis [39]. However, this study included a low number of cases and controls (e.g., 73 cases and 50 controls for the plasma dataset), did not stratify patients for different types of endometriosis, and was not validated in an independent cohort.

Our recent study [51] employed a LR approach, incorporating metabolomics data, the protein biomarker TGFBI, and medical history. Five models, utilizing two metabolite ratios and TGFBI, predicted peritoneal endometriosis with an AUC of 0.889 to 0.914 in the discovery phase and 0.867 to 0.888 in the validation phase. The validated model, which included the ratios of lysoPCa C16:0/Hexoses and lysoPCa C18:2/PCaa C38:3, along with TGFBI, met the criteria for a rule-out triage test. Furthermore, three of the five models met the criteria for a rule-in triage test [51].

### Metabolomics provides novel tools for the diagnosis of endometriosis

The most important result of this study is the identification of metabolomics signatures that can support the early detection of peritoneal endometriosis. This condition is not adequately diagnosed by established gold standard procedures. The use of machine learning techniques has demonstrated that LR [51] and many GLMs can support the diagnosis of peritoneal endometriosis. The present metabolomics-only study showed that patients with different types of endometriosis have distinct GLMs based on metabolic signatures. The present study, although it used the same metabolomic and medical history datasets as LR study, focused on identification of confounder-insensitive signatures composed only of metabolites. Furthermore, we used a different validation strategy in which a single-center population was analyzed first, and then validation was performed in a multicenter population. Our present study and a previous LR study [51] are based on a novel metabolomics dataset, thereby facilitating progress in the use of metabolomics research on endometriosis.

### Physiological aspects of the biomarkers identified in this study

The metabolite ratios and GLMs identified in this study have not been reported as single, diagnostically relevant biomarkers for any other medical indication, nor have they been abandoned as biomarkers. The analysis of individual metabolites contributing to the GLMs aligns with the current understanding of endometriosis pathophysiology. Generally, the essential proteogenic amino acid threonine (Thr) and futher several acylcarnitines (such as C0, C2, C3, C10), which facilitate the fatty acyl shuttle to the mitochondria [92], may reflect the energy demands of proliferating endometriotic lesions. The metabolite C6:1 (hexenoylcarnitine) has been observed to be elevated in obesity [93]. It seems that the same pathway of carnitine metabolism is involved in endometriosis associated metabolome. On the other hand, acylcarnitine C5 indicates enhanced proapoptotic processes [94] and altered valine, isoleucine, and leucine metabolism [95]. Changes in L-arginine (Arg) suggest involvement in nitric oxide synthase signaling [96]. L-proline (Pro) metabolism is linked to glutamine utilization, collagen turnover, and redox regulation [97], which are characteristic of cancer-related collagen matrix remodeling [98]. This might be associated with formation of endometriotic foci and adhesion. L-threonine is also involved in the synthesis of mucin-rich barrier proteins and has been found elevated in type 2 diabetes [99], and chronic kidney disease [100], but lowered in Parkinson’s disease [101]. Changes in the levels of numerous lipids indicate their involvement in the inflammatory processes of endometriosis [43, 44, 102–105]. Lower sphingomyelin profiles, including SM(OH) C22:2, have been previously associated with pneumonia versus COPD exacerbation [106], suggesting a link to epithelial functionality. Phosphatidylcholine levels reflect membrane turnover, lipoprotein transport, and ceramide biosynthesis. These lipids have been controversially discussed in Alzheimer’s biomarker research [107], with PC aa C40:5, lysoPC a C16:0, and SM C18:0 among high-performing discriminating metabolites [108–110]. PC aa 32:0 is considered a hallmark of tumor lipid remodeling [111, 112].. Single lipid species, which constitute the GLMs, have been associated with distinct diseases in this study; however, they have not been used as clinical biomarkers. The GLMs described in this study do not explain the pathogenetic processes of endometriosis, but the models are characteristic of this disease.

### Advantages of this study

This study has several advantages. The cohort size of 513 patients is very large compared to other metabolomic studies, which include e.g. 20-50 patients [46, 102, 113, 114]. Larger studies with e.g. 310 subjects lacked validation and controls were from the healthy women [115]. We introduced a strict definition of entire control group [50] who had the same phenotypes such as pain, dysmenorrhoea, infertility or benign gynecological condition but were not diagnosed with endometriosis by laparoscopy. The use of metabolite ratios rather than single concentrations accounts for the intrinsic variability of the human metabolome [116, 117] and normalises the effects of disease on the metabolome to a degree that supports a validated diagnosis. Machine learning is another advantage as the data were processed in an open and supervised manner. The chosen matrix, the human plasma, can be collected in a semi-invasive way and the SOPs are available worldwide.

### Limitations of the study

We are aware that several important confounding factors such as ethnicity or nutrition were not analysed in this study. The results cannot be generalised to the entire human population as they are based on two center study. In addition, clinical validation [118] of GLMs is required before the biomarkers can be used in precision diagnostics or for therapy monitoring. Such studies require a much larger analysis population including representative numbers of ethnicities, detailed assessment of confounders, a multicenter study and a clinical setting. We further also know that lipids exert their resolving or pro-inflammatory effects depending on the length of the fatty acyl chain and degree of desaturation [119–122]. In-depth analyses of molecular lipid composition in endometriosis diagnostics were beyond the scope of this study.

## Author contributions

A.C. and K.V.: data analysis, interpretation of data; M. N. P.: interpretation of data; A.V.. and R.V.: sample collection, C.P.: experimental work and quality assurance; J.A.: conception, interpretation of data, critical revision, funding, manuscript writing; T.L.R.: study conception and design, critical revision, funding. All authors revised and accepted the final version of the manuscript.

## Institutional Review Board Statement

The study was conducted in accordance with the Declaration of Helsinki and was approved by both national medical ethics committees (Slovenian approval no. 110/05/14, and no. 49/03/16; Austrian approval 545/2010).

## Informed Consent Statement

All participants gave written informed consent before participating in the study.

## Acknowledgements

We thank our study participants who donated their samples and their time. We thank the staff of the Department of Obstetrics and Gynaecology at the University Hospital Ljubljana, Slovenia, especially Tanja Lončar, and at the Medical University of Vienna, especially Manuela Gstöttner, for their support in recruiting study participants. Our special thanks go to Prof. Dr Joško Osredkar and Vera Troha Poljančič from the Medical Centre of the University of Ljubljana, Clinical Institute of Clinical Chemistry and Biochemistry, Medical Centre of the University of Ljubljana, for processing the samples. The authors would also like to thank Julia Scarpa, Dr Werner Römisch-Margl and Katharina Faschinger from Helmholtz Zentrum München, Genome Analysis Centre, Metabolomics Core Facility for sample preparation, measurements and logistics.

## Data sharing

The data underlying this article will be made available by the corresponding author (JA) upon reasonable request, subject to applicable data protection regulations and appropriate safeguards for confidential or personally identifiable information.

## Abbreviations

95%CI: range of confidence interval at 95%
AUC: area under the curve
BMI: body mass index
CV: coefficient of variation
FIA: flow injection
GLM: generalised linear models
HMDB: Human Metabolome Data Bank
LC-ESI-MS/MS: liquid chromatography-electrospray ionisation tandem mass spectrometry
LR: Logistic regression
LLOQ: lower limit of quantification
LOD: limit of detection
NA: not available
NAFLD: non-alcoholic fatty liver disease
MRI: magnetic resonance imaging
OHC: oral hormonal contraception
PLS-R: partial least squares - regression
PLS-DA: partial least squares – discriminant analysis
QC: quality control
RF: random forest
rASRM: revised American Society for Reproductive Medicine
RMSE: root mean square error
RMSEE: root mean square error of estimations
ROC: receiver operator characteristic
SP: specificity
SE: sensitivity
sMRM: scheduled multiple reaction monitoring
MRM: multiple reaction monitoring
SOPs: standard operating procedures

