## Supplementary data for "Metabolomic signatures support the diagnostics of peritoneal endometriosis using generalised linear models"

Supplement Table S1.
Description of metabolites included in GLMs for endometriosis testing

| Metabolite | Trivial name | HMDB ID | CAS number | Chemical Formula |
|---|---|---|---|---|
| PC aa C28:1 | Phosphatidylcholine aa C28:1 | HMDB07867 | na | C36H70NO8P |
| PC ae C30:0 | Phosphatidylcholine ae C30:0 | HMDB13341 | na | C38H78NO7P |
| PC aa C32:0 | Phosphatidylcholine aa C32:0 | HMDB00564 | 63-89-8 | C40H80NO8P |
| PC aa C32:3 | Phosphatidylcholine aa C32:3 | HMDB07876 | na | C40H74NO8P |
| PC aa C34:3 | Phosphatidylcholine aa C34:3 | HMDB08006 | 182820-31-1 | C42H78NO8P |
| PC aa C36:1 | Phosphatidylcholine aa C36:1 | HMDB08037 | na | C44H86NO8P |
| PC aa C36:3 | Phosphatidylcholine aa C36:3 | HMDB07980 | na | C44H82NO8P |
| PC aa C36:4 | Phosphatidylcholine aa C36:4 | HMDB07982 | na | C44H80NO8P |
| PC aa C36:6 | Phosphatidylcholine aa C36:6 | HMDB07892 | na | C44H76NO8P |
| PC aa C38:0 | Phosphatidylcholine aa C38:0 | HMDB07893 | na | C46H92NO8P |
| PC aa C38:3 | Phosphatidylcholine aa C38:3 | HMDB08046 | na | C46H86NO8P |
| PC aa C38:4 | Phosphatidylcholine aa C38:4 | HMDB07988 | na | C46H84NO8P |
| PC aa C40:1 | Phosphatidylcholine aa C40:1 | HMDB13433 | na | C48H96NO7P |
| PC aa C40:4 | Phosphatidylcholine aa C40:4 | HMDB08054 | na | C48H88NO8P |
| PC aa C40:5 | Phosphatidylcholine aa C40:5 | HMDB08055 | na | C48H86NO8P |
| PC aa C42:1 | Phosphatidylcholine aa C42:1 | HMDB08059 | na | C50H98NO8P |
| PC aa C42:2 | Phosphatidylcholine aa C42:2 | HMDB08570 | na | C50H96NO8P |
| PC aa C42:4 | Phosphatidylcholine aa C42:4 | HMDB08572 | na | C50H92NO8P |
| PC aa C42:5 | Phosphatidylcholine aa C42:5 | HMDB08287 | na | C50H90NO8P |
| PC ae C30:2 | Phosphatidylcholine ae C30:2 | HMDB0013410 | na | C38H74NO7P |
| PC ae C32:1 | Phosphatidylcholine ae C32:1 | HMDB0007898 | na | C40H78NO7P |
| PC ae C34:0 | Phosphatidylcholine ae C34:0 | HMDB13405 | na | C42H86NO7P |
| PC ae C34:1 | Phosphatidylcholine ae C34:1 | HMDB0013412 | na | C42H84NO7P |
| PC ae C34:3 | Phosphatidylcholine ae C34:3 | HMDB0013413 | na | C42H80NO7P |
| PC ae C36:0 | Phosphatidylcholine ae C36:0 | HMDB13406 | na | C44H90NO7P |
| PC ae C36:1 | Phosphatidylcholine ae C36:1 | HMDB13427 | na | C44H88NO7P |
| PC ae C36:5 | Phosphatidylcholine ae C36:5 | HMDB11222 | na | C44H78NO7P |
| PC ae C38:0 | Phosphatidylcholine ae C38:0 | HMDB13408 | na | C46H94NO7P |
| PC ae C38:3 | Phosphatidylcholine ae C38:3 | HMDB13439 | na | C46H88NO7P |
| PC ae C38:6 | Phosphatidylcholine ae C38:6 | HMDB13409 | na | C46H82NO7P |
| PC ae C40:0 | Phosphatidylcholine ae C40:0 | HMDB13421 | na | C48H98NO7P |
| PC aa C40:1 | Phosphatidylcholine aa C40:1 | HMDB13433 | na | C48H96NO7P |
| PC ae C40:2 | Phosphatidylcholine ae C40:2 | HMDB13437 | na | C48H96NO7P |
| PC ae C40:3 | Phosphatidylcholine ae C40:3 | HMDB13445 | na | C48H92NO7P |
| PC ae C40:4 | Phosphatidylcholine ae C40:4 | HMDB13442 | na | C48H90NO7P |
| PC ae C40:5 | Phosphatidylcholine ae C40:5 | HMDB13444 | na | C48H88NO7P |
| PC ae C40:6 | Phosphatidylcholine ae C40:6 | HMDB13422 | na | C48H86NO7P |
| PC ae C42:0 | Phosphatidylcholine ae C42:0 | HMDB13443 | na | C50H102NO7P |
| PC ae C42:3 | Phosphatidylcholine ae C42:3 | HMDB13459 | na | C50H96NO7P |
| PC ae C44:3 | Phosphatidylcholine ae C44:3 | HMDB13449 | na | C52H100NO7P |
| PC ae C44:5 | Phosphatidylcholine ae C44:5 | HMDB13456 | na | C52H96NO7P |
| PC ae C44:6 | Phosphatidylcholine ae C44:6 | HMDB13457 | na | C52H94NO7P |
| SM C16:1 | Sphingomyelin C16:1 | HMDB06317 | na | C23H43NO4 |
| SM(OH) C16:1 | Hydroxysphingomyelin C16:1 | HMDB13463 | na | C39H77N2O7P |
| SM(OH) C22:2 | Hydroxysphingomyelin C22:2 | HMDB13467 | na | C45H87N2O7P |
| SM C18:0 | Sphingomyelin C18:0 | HMDB01348 | 58909-84-5 | C41H84N2O6P |
| SM C18:1 | Sphingomyelin C18:1 | HMDB12101 | 108392-10-5 | C41H81N2O6P |
| SM C20:2 | Sphingomyelin 20:2 | HMDB13465 | na | C43H83N2O6P |
| SM C22:3 | Sphingomyelin C22:3 | HMDB13468 | na | C45H85N2O6P |

| Metabolite | Trivial name | HMDB ID | CAS number | Chemical Formula |
|---|---|---|---|---|
| SM C24:1 | Sphingomyelin C24:1 | HMDB12107 | 94359-13-4 | C47H93N2O6P |
| lysoPC a C14:0 | Lysophosphatidylcholine a C14:0 | HMDB10379 | 20559-16-4 | C22H46NO7P |
| lysoPC a C16:0 | Lysophosphatidylcholine a C16:0 | HMDB10382 | 17364-16-8 | C24H50NO7P |
| LysoPC a C17:0 | Lysophosphatidylcholine a C17:0 | HMDB12108 | 50930-23-9 | C25H52NO7P |
| lysoPC a C18:1 | Lysophosphatidylcholine a C18:1 | HMDB02815 | 19420-56-5 | C26H52NO7P |
| lysoPC a C18:2 | Lysophosphatidylcholine a C18:2 | HMDB10386 | 22252-07-9 | C26H50NO7P |
| lysoPC a C20:4 | Lysophosphatidylcholine a C20:4 | HMDB10395 | 60701-99-7 | C28H50NO7P |
| C0 | L-Carnitine (free carnitine) | HMDB00062 | 541-15-1 | C7H15NO3 |
| C3 | Propionylcarnitine | HMDB00824 | 20064-19-1 | C10H20NO4 |
| C3-DC (C4-OH) | Hydroxybutyrylcarnitine | HMDB02095 | 910825-21-7 | C10H17NO6 |
| C4 | Isobutyryl-L-carnitine | HMDB00736 | 25518-49-4 | C11H21NO4 |
| C5 | Isovalerylcarnitine | HMDB00688 | 31023-24-2 | C12H23NO4 |
| C5:1 | Tiglylcarnitine | HMDB02366 | 64681-36-3 | C12H21NO4 |
| C5-M-DC | Methylglutaryl-L-carnitine | HMDB00552 | 102673-95-0 | C12H25NO5 |
| C6:1 | Hexenoylcarnitine | HMDB13161 | na | C13H23NO4 |
| C8 | Octanoylcarnitine | HMDB0000791 | 25243-95-2 | C15H30NO4 |
| C10 | Decanoylcarnitine | HMDB00651 | 1492-27-9 | C17H33NO4 |
| C10:1 | Decenoylcarnitine | HMDB13205 | na | C17H31NO4 |
| C12-DC | Dodecanedioylcarnitine | HMDB13327 | na | C19H35NO6 |
| C14:1 | Tetradecenoylcarnitine | HMDB02014 | 835598-21-5 | C21H39NO4 |
| C14:2 | Tetradecadienylcarnitine | HMDB13331 | na | C21H37NO4 |
| C14:2-OH | Hydroxytetradecadienylcarnitine | HMDB240755 | na | C21H37NO5 |
| C16 | Hexadecanoylcarnitine | HMDB00222 | 2364-67-2 | C23H45NO4 |
| C16:2-OH | Hydroxyhexadecadienylcarnitine | HMDB13335 | na | C23H41NO5 |
| C18 | Stearoylcarnitine | HMDB00848 | 25597-09-5 | C25H50NO4 |
| C18:2 | Octadecadienylcarnitine | HMDB06461 | 85114-47-2 | C25H45NO4 |
| Arg | L-Arginine | HMDB00517 | 74-79-3 | C6H14N4O2 |
| Gln | L-Glutamine | HMDB00641 | 56-85-9 | C5H10N2O3 |
| Gly | L-Glycine | HMDB00123 | 56-40-6 | C2H5NO2 |
| Met | Methionine | HMDB00696 | 63-68-3 | C5H11NO2S |
| Orn | L-Ornitine | HMDB00214 | 3184-13-2 | C5H12N2O2 |
| Pro | L-Proline | HMDB00162 | 147-85-3 | C5H9NO2 |
| Ser | L-Serine | HMDB00187 | 56-45-1 | C3H7NO3 |
| Thr | L-Threonine | HMDB00167 | 72-19-5 | C4H9NO3 |
| Trp | L-Tryptophan | HMDB00929 | 73-22-3 | C11H12N2O2 |
| Tyr | L-Tyrosine | HMDB00158 | 60-18-4 | C9H11NO3 |
| SFA_PC | composite lipid class: saturated glycerophosphocholines | na | na | na |

Abbreviations are provides as described the AbsoluteIDQ™ p180 assay [58]. Acylcarnitines (e.g. C6:1) are given as x:y (placeholders x and y represent the total number of carbons and double bonds of all chains, respectively), amino acids abbreviated in three letter code (e.g. Gln), phosphatidylcholines as PC nn x:y (nn denotes either "aa" for diacyl, "ae" for acyl-alkyl), lysophosphatidylcholines as lysoPC a x:y, ("a" denotes acyl) , and sphingolipids as SM x:y, ", "na" – not annotated in HMDB (human Metabolome Data Bank) [59]. Amino acids are abbreviated in three letter code. Chemical names of metabolites are given without a resolution of isobars. The names of individual metabolites are defined as state-of-the-art in the research area [43, 44, 60].

Supplement Table S2

Demographic description and clinical characteristics of discovery and validation population.

Discovery phase of the study

| Variable | Cases n=151 | Cases with peritoneal endometriosis n=52 | Controls n=84 | p-value [1] |
|---|---|---|---|---|
| Age (years) | | | | |
| Mean (Standard deviation) [Min, Max] | 30.56 (4.1) [18, 41] | 30.44 (3.65) [22, 37] | 30.35 (4.53) [20, 38] | 0.716 [2] 0.897 [3] |
| BMI (kg/m$^2$) | | | | |
| Mean (Standard deviation) [Min, Max] | 22.8 (3.75) [16, 37.3] | 22.51 (3.1) [16, 32] | 24.27 (4.96) [18.3, 41.9] | 0.011 [2] 0.023 [3] |
| Menstrual cycle phase, n (%) | | | | |
| Proliferative | 73 (48.34) | 27 (51.92) | 37 (44.05) | 0.388 [2] |
| Secretory | 60 (39.74) | 24 (46.15) | 34 (40.48) | 0.074 [3] |
| Oral hormonal contraception | 17 (11.26) | 1 (1.92) | 11 (13.1) | |
| Irregular or no cycle | 0 | 0 | 2 (2.38) | |
| Missing | 1 (0.66) | 0 | 0 | |
| Regular menstrual cycle, n (%) | | | | |
| Yes | 138 (91.39) | 51 (98.08) | 67 (79.76) | 0.012 [2] |
| No | 10 (6.62) | 1 (1.92) | 17 (20.24) | 0.001 [3] |
| Missing | 3 (1.99) | 0 | 0 | |
| Oral contraception in the last 3 months before the surgery, n (%) | | | | |
| Yes | 9 (5.96) | 2 (3.85) | 6 (7.14) | 0.861 [2] |
| No | 141 (93.38) | 50 (96.15) | 78 (92.86) | 0.710 [3] |
| Missing | 1 (0.66) | 0 | 0 | |
| Hormonal therapy in the last 3 months before the surgery, n (%) | | | | |
| Yes | 22 (14.57) | 9 (17.31) | 16 (19.05) | 0.653 [2] |
| No | 128 (84.77) | 43 (82.69) | 68 (80.95) | 1 [3] |
| Missing | 1 (0.66) | 0 | 0 | |
| Medications in the last week before the surgery, n (%) | | | | |
| Yes | 52 (34.44) | 18 (34.62) | 20 (23.81) | 0.172 [2] |
| No | 97 (64.24) | 33 (63.46) | 62 (73.81) | 0.348 [3] |
| Missing | 2 (1.32) | 1 (1.92) | 2 (2.38) | |

| | | | | |
|---|---|---|---|---|
| Type of endometriosis, n (%) | | | | |
| Ovarian | 25 (16.56) | NA | NA | |
| Peritoneal | 52 (34.44) | 52 (100) | NA | |
| Ovarian and peritoneal | 42 (27.81) | NA | NA | |
| Deep infiltrating | 4 (2.65) | NA | NA | |
| Ovarian and deep | 9 (5.96) | NA | NA | |
| Peritoneal and deep i | 5 (3.31) | NA | NA | |
| Ovarian, peritoneal and deep | 14 (9.27) | NA | NA | |
| Missing | 0 | NA | NA | |
| rAFS stage, n (%) | | | | |
| I | 52 (34.44) | 48 (92.31) | NA | |
| II | 15 (9.93) | 4 (7.69) | NA | |
| III | 56 (37.09) | 0 | NA | |
| IV | 24 (15.89) | 0 | NA | |
| Missing | 4 (2.65) | 0 | NA | |
| Alcohol consumption, n (%) | | | | |
| Never | 43 (28.48) | 16 (30.77) | 25 (29.76) | 0.788 [2] |
| Occasionally | 89 (58.94) | 31 (59.62) | 54 (64.29) | 0.771 [3] |
| Once a week | 11 (7.28) | 3 (5.77) | 4 (4.76) | |
| 2 to 3 times per week | 6 (3.97) | 2 (3.85) | 1 (1.19) | |
| More than 3 times per week | 1 (0.66) | 0 | 0 | |
| Missing | 1 (0.66) | 0 | 0 | |
| Smoking status, n (%) | | | | |
| Non smoker | 106 (70.2) | 35 (67.31) | 54 (64.29) | 0.354 [2] |
| Smoker | 28 (18.54) | 9 (17.31) | 19 (22.62) | 0.743 [3] |
| Occasional (once weekly) | 7 (4.64) | 3 (5.77) | 3 (3.57) | ] |
| Occasional (once monthly) | 1 (0.66) | 1 (1.92) | 4 ( 4.76) | |
| Former smoker | 8 (5.3) | 4 (7.69) | 4 (4.76) | |
| Missing | 1 (0.66) | 0 | 0 | |
| Sport or recreation 2 days before the surgery, n (%) | | | | |
| Yes | 44 (29.14) | 21 (40.38) | 22 (26.19) | 0.646 [2] |
| No | 105 (69.54) | 31 (59.62) | 62 (73.81) | 0.091 [3] |
| Missing | 2 (1.32) | 0 | 0 | |

Validation phase of the study

| Variable | Cases n=166 | Cases with peritoneal endometriosis n=42 | Controls n=112 | p-value [1] |
|---|---|---|---|---|
| Age (years) | | | | |
| Mean (Standard deviation) [Min, Max] | 32.15 (6.09) [19.9, 50.5] | 29.9 (4.54) [19.9, 38.2] | 34.1 (8.22) [18.1, 54.1] | 0.024 [2] 0.002 [3] |
| BMI (kg/m$^2$) | | | | |
| Mean (Standard deviation) [Min, Max] | 22.86 (4.46) [16.9, 50.1] | 22.88 (3.85) [18.2, 34] | 24.23 (4.63) [17.6, 42.2] | 0.014 [2] 0.092 [3] |
| Menstrual cycle phase, n (%) | | | | |
| Proliferative | 88 (53.01) | 21 (50) | 48 (42.86) | 0.009 [2] |
| Secretory | 67 (40.36) | 20 (47.62) | 42 (37.5) | 0.088 [3] |
| Oral hormonal contraception | 6 (3.61) | 1 (2.38) | 6 (5.36) | |
| Irregular or no cycle | 1 (0.6) | 0 | 4 (3.57) | |
| Missing | 4 (2.41) | 0 | 12 (10.71) | |
| Regular menstrual cycle, n (%) | | | | |
| Yes | 145 (87.35) | 37 (88.1) | 79 (70.54) | 0.002 [2] |
| No | 17 (10.24) | 5 (11.9) | 27 (24.11) | 0.06 [3] |
| Missing | 4 (2.41) | 0 | 6 (5.36) | |
| Oral contraception in the last 3 months before the surgery, n (%) | | | | |
| Yes | 46 (27.71) | 5 (11.9) | 21 (18.75) | 0.184 [2] |
| No | 118 (71.08) | 37 (88.1) | 89 (79.46) | 0.441 [3] |
| Missing | 2 (1.2) | 0 | 2 (1.79) | |
| Hormonal therapy in the last 3 months before the surgery, n (%) | | | | |
| Yes | 21 (12.65) | 5 (11.9) | 12 (10.71) | 0.797 [2] |
| No | 143 (86.14) | 37 (88.1) | 98 (87.5) | 1 [3] |
| Missing | 2 (1.2) | 0 | 2 (1.79) | |
| Medications in the last week before the surgery, n (%) | | | | |
| Yes | 74 (44.58) | 12 (28.57) | 56 (50) | 0.554 [2] |
| No | 90 (54.22) | 30 (71.43) | 54 (48.21) | 0.026 [3] |
| Missing | 2 (1.2) | 0 | 2 (1.79) | |
| Type of endometriosis, n (%) | | | | |

| | | | | |
|---|---|---|---|---|
| Ovarian | 32 (19.28) | NA | NA | |
| Peritoneal | 42 (25.3) | 42 (100) | NA | |
| Ovarian and peritoneal | 39 (23.49) | NA | NA | |
| Deep infiltrating | 9 (5.42) | NA | NA | |
| Ovarian and deep | 16 (9.64) | NA | NA | |
| Peritoneal and deep | 9 (5.42) | NA | NA | |
| Ovarian, peritoneal and deep | 18 (10.84) | NA | NA | |
| Missing | 1 (0.6) | NA | NA | |
| rAFS stage, n (%) | | | | |
| I | 42 (25.3) | 34 (80.95) | NA | |
| II | 38 (22.89) | 8 (19.05) | NA | |
| III | 49 (29.52) | 0 | NA | |
| IV | 31 (18.67) | 0 | NA | |
| Missing | 6 (3.61) | 0 | NA | |
| Alcohol consumption, n (%) | | | | |
| Never | 39 (23.49) | 13 (30.95) | 22 (19.64) | 0.101 [2] |
| Occasionally | 106 (63.86) | 24 (57.14) | 61 (54.46) | 0.216 [3] |
| Once a week | 9 (5.42) | 1 (2.38) | 16 (14.29) | |
| 2 to 3 times per week | 7 (4.22) | 2 (4.76) | 8 (7.14) | |
| More than 3 times per week | 2 (1.2) | 1 (2.38) | 2 (1.79) | |
| Missing | 3 (1.81) | 1 (2.38) | 3 (2.68) | |
| Smoking status, n (%) | | | | |
| Non smoker | 99 (59.64) | 19 (45.24) | 52 (46.43) | 0.117 [2] |
| Smoker | 46 (27.71) | 15 (35.71) | 41 (36.61) | 0.565 [3] |
| Occasional (once weekly) | 2 (1.2) | 0 | 4 (3.57) | |
| Occasional (once monthly) | 3 (1.81) | 1 (2.38) | 0 | |
| Former smoker | 13 (7.83) | 6 (14.29) | 13 (11.61) | |
| Missing | 3 (1.81) | 1 (2.38) | 2 (1.79) | |
| Sport or recreation 2 days before the surgery, n (%) | | | | |
| Yes | 47 (28.31) | 16 (38.1) | 19 (16.96) | 0.072 [2] |
| No | 116 (69.88) | 25 (59.52) | 89 (79.46) | 0.018 [3] |
| Missing | 3 (1.81) | 1 (2.38) | 4 (3.57) | |

BMI = Body mass index; rAFS = revised American Fertility Society stage; Min = minimum; Max = Maximum; NA = Not applicable

[1] Two sample t-test was used for testing continuous and Fisher's exact test was used for testing categorical variables.

[2] Test was performed for comparing all case vs. all control participants.

[3] Test was performed for comparing peritoneal endometriosis vs. all control participants.

Supplement Table S3. Listing of additional GLMs for different types of endometriosis in discovery population.

| # | GLM all type of endometriosis | AUC | RMSE | SP % | SE % |
|---|---|---|---|---|---|
| 2 | Arg/PC ae C36:0 + lysoPC a C16:0/SM C18:1 + PC ae C38:0/PC ae C40:0 | 0.72 | 0.0434 | 87 | 73 |
| 3 | Thr/PC aa C34:3 + lysoPC a C17:0/SM (OH) C16.1 + Arg/PC ae C36:0 | 0.72 | 0.0766 | 88 | 71 |
| 4 | Thr/PC aa C34:3 + Arg/PC ae C36:0 + lysoPC a C16:0/SM C18:1 | 0.71 | 0.0713 | 87 | 71 |
| 5 | Arg/PC aa C36:6 + lysoPC a C17:0/SM (OH) C16:1 + Arg/PC ae C36:0 | 0.71 | 0.0544 | 86 | 71 |
| 6 | lysoPC a C17:0/SM (OH) C16:1 + Arg/PC ae C36:0 + C18/lysoPC a C14:0 | 0.71 | 0.0540 | 88 | 62 |
| # | GLM peritoneal mixed | AUC | RMSE | SP % | SE % |
| 2 | Arg/PC aa C36:6 + C5/lysoPC a C17:0 + C5/Arg | 0.68 | 0.0860 | 73 | 63 |
| 3 | Orn/PC ae C38:0 + C5/lysoPC.a.C17.0 + C5/Arg | 0.67 | 0.0821 | 71 | 65 |
| 4 | C0/Gly + Orn/PC ae C38:0 + Tyr/PC aa C42:2 | 0.67 | 0.0648 | 67 | 64 |
| 5 | SM C18:0 + C5/lysoPC a C17:0 + C5/Arg | 0.67 | 0.0955 | 69 | 63 |
| 6 | C5/lysoPC a C17:0 + C5/Arg + Ser/SM(OH) C16:1 | 0.67 | 0.0842 | 68 | 63 |
| # | GLM ovarian | AUC | RMSE | SP % | SE % |
| 2 | PC aa C38:0/PC ae C36:1 + Thr/SM (OH) C22:1 + lysoPC a C14:0/PC aa C28:1 | 0.70 | 0.0782 | 77 | 73 |
| 3 | Thr/SM (OH) C22:1 + PC aa C28:1/PC ae C34:3 + C18:2/ PC ae C34:3 | 0.70 | 0.0982 | 92 | 80 |
| 4 | PC aa C36:3/PC ae C40:5 + C3/PC ae C34:1 + Met/PC aa C36:3 | 0.70 | 0.0893 | 97 | 78 |
| 5 | PC aa C36:3/PC ae C40:5 + PC aa C28:1/PC ae C34:3 + Gly/PC ae C36:1 | 0.70 | 0.0414 | 97 | 77 |
| 6 | PC aa C38:0/PC ae C36:1 + C18:2/PC ae C34:3 + Met/PC aa C36:3 | 0.69 | 0.1026 | 73 | 80 |

| # | GLM ovarian mixed | AUC | RMSE | SP % | SE % |
|---|---|---|---|---|---|
| | C10/PC aa C36:6 + Pro/PC ae C34:0 + PC ae C42:3/SM (OH) C16:1 | 0.67 | 0.0716 | 63 | 60 |
| | C10/PC aa C36:6 + PC ae C42:3/SM (OH) C16:1 + C6:1/lysoPC a C20:4 | 0.66 | 0.0688 | 61 | 61 |
| | C10/PC aa C36:6 + PC ae C42:3/SM (OH) C16:1 + lysoPC a C20:4/PC ae C40:2 | 0.66 | 0.0709 | 61 | 60 |
| | Ser/PC aa C38:3 + C10/PC aa C36:6 + PC ae C42:3/SM (OH) C16:1 | 0.66 | 0.0789 | 58 | 58 |
| | C10/lysoPC a C18:1 + C10/PC aa C36:6 + PC ae C42:3/SM (OH) C16:1 | 0.66 | 0.1236 | 63 | 63 |

Abbreviations: metabolite abbreviations are explained in supplementary Table S1, AUC – area under the curve, RMSE - root mean square error, SP – specificity, SE - sensitivity.

Please note that the corresponding peritoneal endometriosis data were displayed in the table 1. Only further GLMs for each type of endometriosis are provided. The best GLM for each type of endometriosis was already presented in Table 2.

Supplement Figure S1. Cross validation of different GLMs for the peritoneal endometriosis in the discovery population.

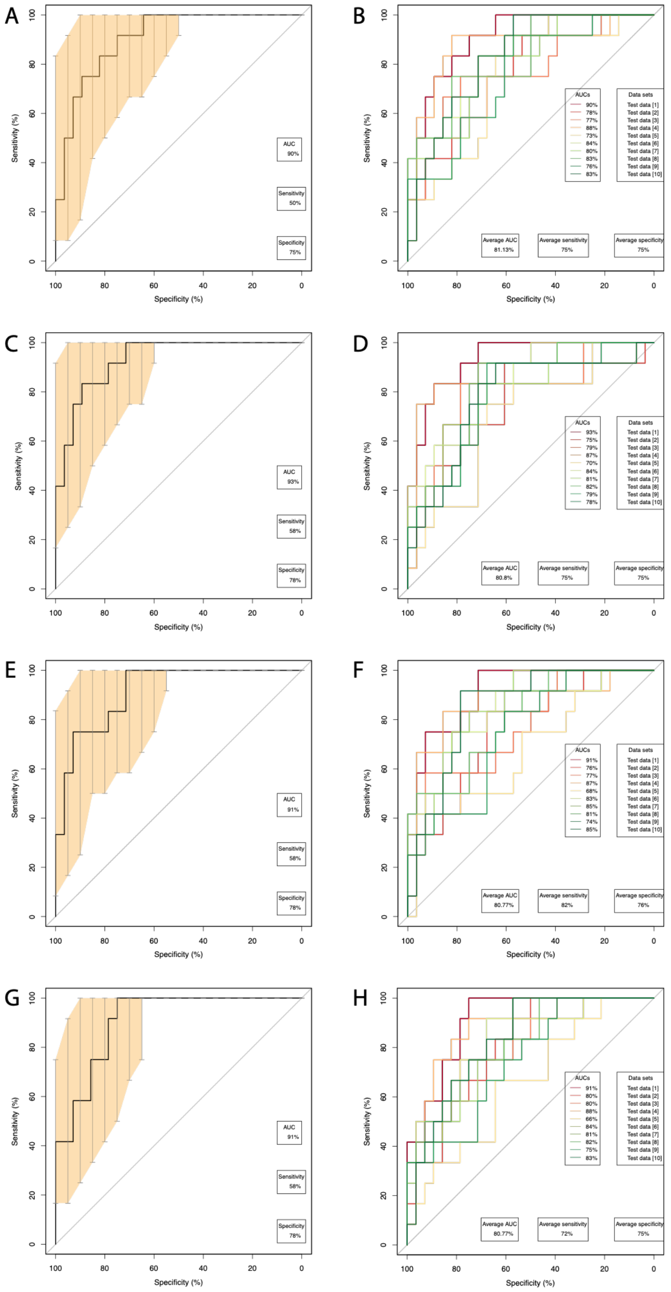

A) Best cross validated GLM for PC aa C40:5/SFA_ PC + lysoPC a C16:0/SM(OH) C16: 1 + PC aa C32:0/SM C18:0. The light orange area and the bars depict 95%CI.
B) Results of 10-fold cross validation and the average AUC, specificity and sensitivity for the model in A.
C) Best cross validated GLM for lysoPC a C16:0/SM(OH) C16:1 + PC aa C32:0/SM C18:0 + PC aa C32:0/PC aa C38:3. The light orange area and the bars depict 95%CI.
D) Results of 10-fold cross validation and the average AUC, specificity and sensitivity for the model in C.
E) Best cross validated GLM for C6:1_div_by_Pro/PC aa C40:5/SFA_PC + lysoPC a C16:0/SM(OH) C16:1. The light orange area and the bars depict 95%CI.
F) Results of 10-fold cross validation and the average AUC, specificity and sensitivity for the model in E.
G) Best cross validated GLM for PC aa C40:5/SFA_PC + lysoPC a C16:0_div_by_SM(OH) C16:1 + Arg/PC ae C34:0. The light orange area and the bars depict 95%CI.
H) Results of 10-fold cross validation and the average AUC, specificity and sensitivity for the model in G.

Supplementary Figure S2. Cross validation of different GLMs for the different types of endometriosis in the discovery population.

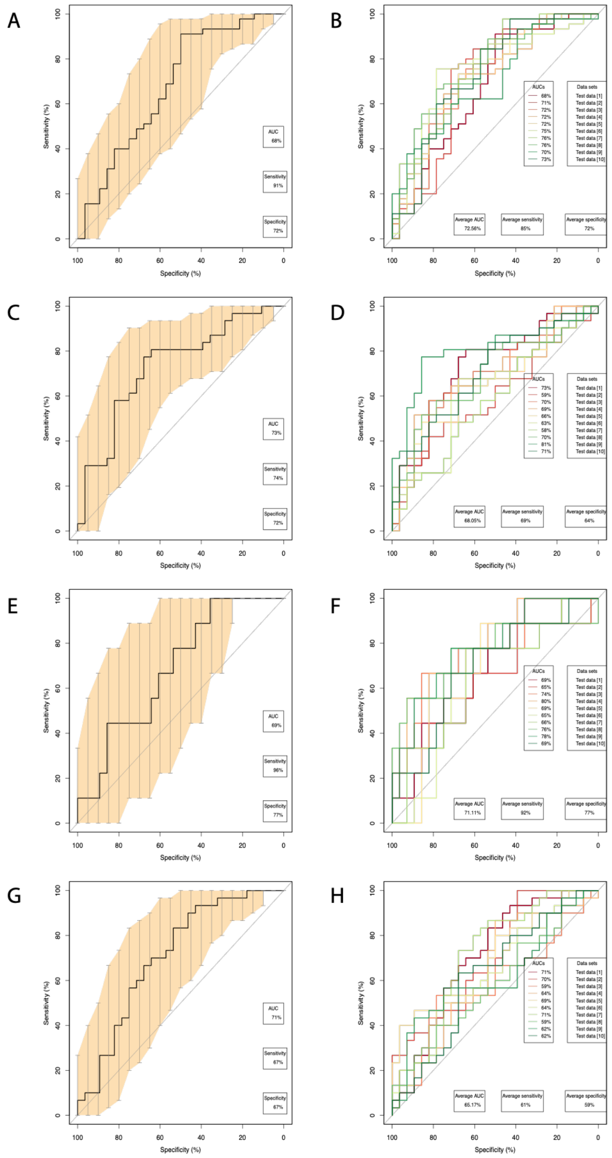

A) Best model for all types of endometriosis: LysoPC a C17:0/SM(OH) C16:1 + Arg/PC ae C36:0 + PC ae C38:0/PC ae C40:0
B) Results of 10-fold cross validation and the average AUC, specificity and sensitivity for the model in A for all types of endometriosis
C) Best model for peritoneal mixed endometriosis: Orn/PC ae C38:0 + C4/PC aa C38:4 + Tyr/PC aa C42:2
D) Results of 10-fold cross validation and the average AUC, specificity and sensitivity for the model in C for peritoneal mixed endometriosis
E) best model for ovarian endometriosis: PC aa C36:3/PC ae C40:5 + lysoPC a C14:0/PC aa C28:1 + Met/PC aa C36:3
F) Results of 10-fold cross validation and the average AUC, specificity and sensitivity for the model in E for ovarian endometriosis
G) best model for ovarian mixed endometriosis :C10/PC aa C36:6 + Pro/PC ae C34:0 + PC ae C42:3/SM(OH) C16:1
H) Results of 10-fold cross validation and the average AUC, specificity and sensitivity for the model in G for ovarian mixed endometriosis

Please note that the GLMs for peritoneal endometriosis is not included as it has been shown already in Figure 5.
